# Phylogenetic tree inference using generative models

**DOI:** 10.64898/2026.06.14.732140

**Authors:** Edo Dotan, Asaf Schers, Elya Wygoda, Tal Pupko, Yonatan Belinkov

## Abstract

Accurate inference of phylogenetic trees is fundamental to evolutionary biology, yet existing methods rely on complex pipelines involving multiple sequence alignment, explicit evolutionary models, and computationally intensive tree search procedures. Here, we present BetaInfer, a generative framework that reformulates phylogenetic tree inference as a sequence transduction problem. BetaInfer leverages hybrid transformer-based architectures to directly map sets of unaligned sequences to phylogenetic trees represented in Newick format. Trained on large-scale simulated evolutionary data with known ground truth, BetaInfer learns to capture complex evolutionary signals directly from sequence data. Ensemble-based generation of multiple candidate trees further improves robustness, reducing reconstruction error by over 30% relative to single predictions. Across extensive evaluations on both simulated and empirical datasets, BetaInfer achieves competitive performance relative to state-of-the-art phylogenetic pipelines, matching, and in some cases exceeding, the accuracy of established likelihood-based and distance-based methods under a wide range of conditions. Interpretability analyses reveal that BetaInfer leverages internal pairwise-distance computations to synthesize evolutionary relationships into an integrated, global representation that supports direct tree generation. Together, these results demonstrate that generative models can serve as a viable and scalable alternative to standard phylogenetic pipelines.

## Introduction

Phylogenetic tree inference is a cornerstone of evolutionary biology, providing a framework for reconstructing evolutionary relationships among species or genes and tracing patterns of divergence (Felsenstein, 2004; Yang & Rannala, 2012). This conceptual framework traces back to Darwin’s theory of common descent, which first articulated the idea that all species are connected through a branching evolutionary history (Darwin, 1859). Accurate phylogenetic reconstruction underpins a wide range of biological analyses, including the study of adaptation (Mi et al., 2016), speciation (Igea & Tanentzap, 2020), and the timing of evolutionary events (Gavryushkina et al., 2017). Errors in inferred tree topologies or branch lengths can propagate to downstream analyses, leading to incorrect inferences about ancestral states, rates of evolution, and patterns of trait evolution. As phylogenetic trees increasingly serve as foundational inputs for large-scale genomic and evolutionary studies, ensuring their accuracy is essential for drawing reliable biological inferences across diverse datasets and evolutionary timescales (Lemmon & Moriarty, 2004; Warnow, 2012).

Distance-based phylogenetic inference methods from sequence data first transform the data into a matrix of pairwise distances between sequences and then infer a tree directly from these distances. In the Unweighted Pair Group Method with Arithmetic Mean (UPGMA), the tree is inferred by iteratively grouping taxa (or genes) according to their estimated evolutionary distances, offering computational efficiency and conceptual simplicity. UPGMA assumes a molecular clock and produces ultrametric trees (Sokal & Michener, 1958), an assumption that is often violated in real datasets (Drummond et al., 2006; Flouri et al., 2022). The neighbor-joining (NJ) method relaxes the clock assumption and can be viewed as a heuristic for estimating trees under the minimum evolution criterion (Saitou & Nei, 1987). Distance-based approaches often estimate distances from all pairwise alignments among the analyzed sequences and therefore do not require multiple sequence alignments (MSAs). However, information regarding positional homology is often lost when reducing sequence data to a distance matrix (Felsenstein, 2004).

Maximum parsimony is a character-based approach that seeks phylogenetic trees minimizing the total number of evolutionary changes required to explain the observed sequences (Fitch, 1971; Kluge & Farris, 1969). It does not rely on an explicit probabilistic model of sequence evolution. In the standard Wagner parsimony framework, all substitutions are weighted equally, an assumption that is often unrealistic. In addition, variation in evolutionary rates among sites is not accounted for, further limiting parsimony’s ability to capture the sequence dynamics underlying biological data. These simplifying assumptions have been shown to reduce the accuracy of parsimony relative to more advanced approaches, such as maximum likelihood and Bayesian inference (Hall, 2005; Kuhner & Felsenstein, 1994).

Modern phylogenetic inference methods assume an explicit probabilistic model of sequence evolution. This model specifies substitution probabilities as well as how evolutionary rates vary across sites. Similar to parsimony, these methods do not operate directly on raw sequence data, but instead require an MSA that defines positional homology across sequences (Notredame et al., 2000; Thompson et al., 1994). Given an MSA, a tree topology with branch lengths, and a stochastic model of sequence evolution, the likelihood of the observed data can be efficiently computed. Maximum likelihood inference entails a combinatorial search over tree topologies and branch lengths to identify the tree that maximizes this likelihood (Felsenstein, 1981; Yang, 1994). In Bayesian inference, a posterior distribution is estimated over trees and model parameters (Drummond & Rambaut, 2007; Huelsenbeck & Ronquist, 2001; Yang & Rannala, 1997).

Despite the improvement in inference accuracy achieved by the shift from parsimony and distance-based methods to probabilistic approaches, challenges in tree inference remain. For example, evolutionary dynamics vary substantially among genes and lineages. Substitution patterns also vary across different structural regions of proteins and among non-coding genomic elements, such as introns, promoters, and enhancers. They further depend on genomic context, including whether genes are encoded in the nucleus, mitochondria, or plastids (Abascal et al., 2007). The ability of probabilistic models to capture such heterogeneity remains limited. Moreover, most models do not explicitly account for indel dynamics, which also vary considerably among phylogenetic groups (Loewenthal et al., 2021; Löytynoja & Goldman, 2008; Wygoda et al., 2024). These models typically assume conditional independence among sites to ensure computational feasibility; however, this assumption often fails to capture the complexity of real evolutionary processes, including lineage-specific effects and site dependencies (Redelings et al., 2024; Thorne, 2000), and can introduce systematic biases in estimates of tree topologies and branch lengths. In addition to model misspecification, inference errors can also arise from inaccuracies in the MSA and from inefficient exploration of tree space (Philippe et al., 2011; Warnow, 2012).

Machine learning has recently been introduced to improve phylogenetic inference (reviewed in Buch & Gambhava, 2026; Mo et al., 2024). Some approaches use learned heuristics to guide tree search, training models to predict likely neighboring trees and thereby accelerating the search without explicit likelihood computation (Azouri et al., 2021). Other work has proposed training deep learning classifiers to directly predict tree topologies from MSAs (Zou et al., 2020). However, because the number of possible tree topologies grows super-exponentially with the number of taxa, such classifiers are inherently limited to small taxon sets, typically quartets or quintets. Recent end-to-end deep learning frameworks integrate learned distance-based methods with deep sequence encoding (Nesterenko et al., 2025; Zhang et al., 2025), illustrating a promising direction toward machine learning–assisted phylogenetic inference that balances accuracy and computational feasibility.

Recent advances in deep learning, particularly those inspired by natural language processing (NLP), have provided new opportunities for modeling the complex dependencies that characterize biological sequences (Alley et al., 2019; Leclercq & Droit, 2025; Rannon & Burstein, 2025; Rao et al., 2021; Rives et al., 2021). In particular, transformer-based architectures have demonstrated strong performance across a wide range of sequence-based tasks, including prediction of protein structure (Jumper et al., 2021), stability (Umerenkov et al., 2023), and localization and functional annotation (Elnaggar et al., 2022). These deep learning models learn rich contextual representations that implicitly encode evolutionary constraints and long-range interactions, making them well suited for downstream evolutionary and comparative analyses.

Here, we introduce BetaInfer, a deep learning framework for phylogenetic tree inference. Our approach builds on a hybrid state-space model (SSM)–transformer architecture trained on simulated evolutionary data. To the best of our knowledge, this represents the first application of hybrid transformer architectures to phylogenetic inference. Rather than relying on predefined MSAs, fixed tree-search heuristics, or explicit parametric models of sequence evolution, BetaInfer learns to reconstruct phylogenetic relationships directly from unaligned sequence data. This design provides substantial flexibility, enabling the method to accommodate diverse datasets and complex evolutionary regimes. Our results indicate that generative models offer a compelling alternative to conventional phylogenetic inference pipelines. More broadly, this work highlights the potential of deep learning to reshape phylogenetic analysis by enabling scalable, end-to-end, data-driven reconstruction of evolutionary histories.

## New approaches

### The generative algorithm for inferring phylogenetic trees

Our hybrid transformer-based model for phylogenetic tree inference is trained using a next-token prediction objective. In this framework, the model learns to predict each subsequent token conditioned on all preceding tokens. This paradigm has a long-standing history in language modeling, demonstrating that sequential prediction is highly effective at capturing complex dependencies in discrete sequences (Bengio et al., 2003; Mikolov et al., 2010; Radford, et al., 2019). Next-token prediction is now the core training objective of modern text and code generation systems (Dubey et al., 2024; Gemini Team et al., 2024; OpenAI et al., 2024). In other words, the model learns to complete sequences by leveraging contextual information encoded in earlier tokens. Tokens constitute the fundamental units that map discrete symbols to numerical representations processed by the model (Dotan et al., 2024; Gastaldi et al., 2025; Mielke et al., 2021).

A collection of evolutionarily related, unaligned sequences does not naturally correspond to a single “sentence,” the standard input format for next-token prediction tasks. Therefore, a critical preprocessing step is to transform these sequences, which constitute the input to our phylogenetic tree reconstruction task, into a unified sequential representation (Fig. 1). Among several possible encoding strategies (Dotan et al., 2023), we adopt a “concat” representation, in which unaligned sequences are concatenated into a single input sequence using a dedicated separator token (the pipe symbol, “|”). At the end of this concatenated input, we append a special token that signals the start of phylogenetic tree inference (e.g., “<INFER>”; Fig. 1). Following this token, the model is tasked with generating a representation of the phylogenetic tree in Newick format (Czech et al., 2017). Put simply, the input to the model consists of concatenated unaligned sequences followed by the <INFER> token, and the output is a Newick string representing the tree topology and branch lengths. Notably, the Newick representation of a tree is not unique: although there is a single underlying tree topology for a given set of sequences, multiple valid Newick strings may encode the same tree topology and branch lengths.

**Figure 1.**
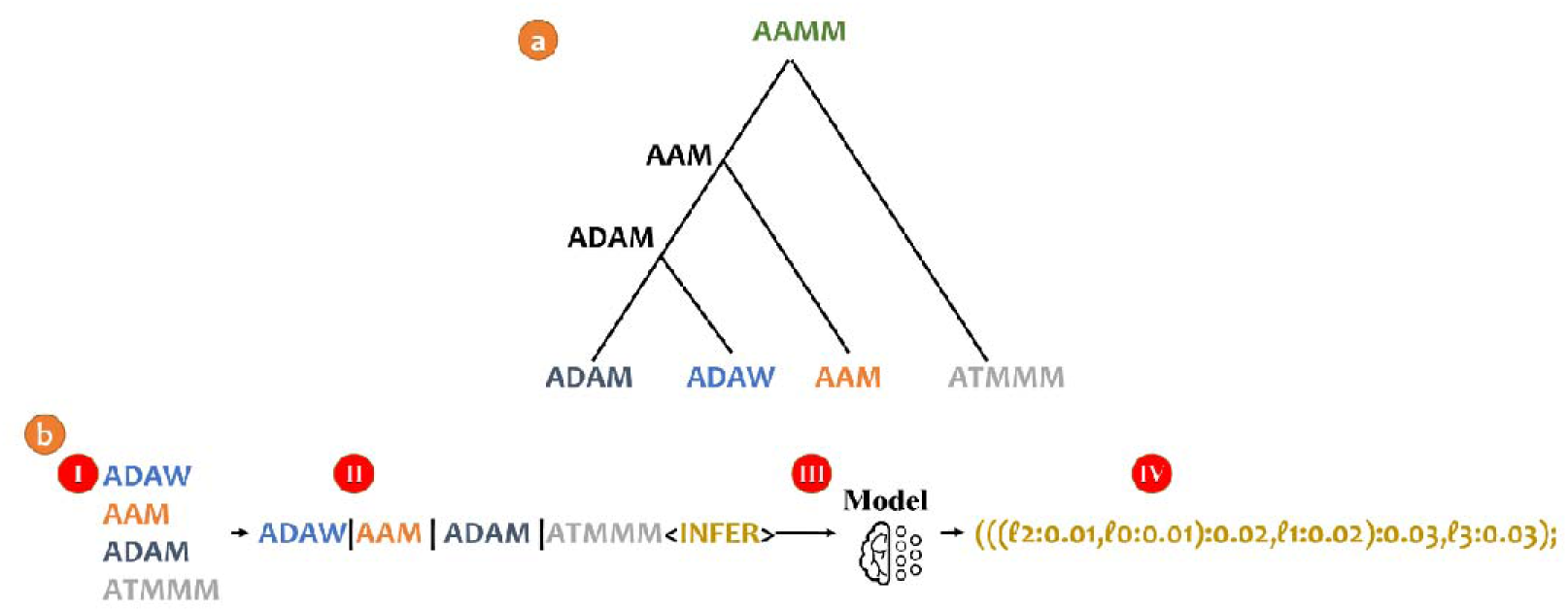
Illustration of phylogenetic tree prediction using BetaInfer. Panel (a) shows the simulated “true” evolutionary dynamics, in which the ancestral sequence “AAMM” diverges along a phylogenetic tree into four leaves: “ADAM”, “ADAW”, “AAM”, and “ATMMM”. Panel (b) illustrates the BetaReconstruct pipeline: (I) unaligned protein sequences serve as input to the model; (II) the sequences are concatenated into a single input “sentence”, and a special token marking the start of phylogenetic inference is appended; (III) the trained model processes the input; and (IV) the model generates the output phylogenetic tree. Sequence identifiers are not provided as input. In the output tree, the label ℓ₀ corresponds to the first concatenated sequence, ℓ₁ to the second, and so on.

In contrast to classical phylogenetic pipelines, our generative formulation does not require an explicit evolutionary model, an MSA, or a predefined tree-search algorithm at the inference stage (an evolutionary model is used to generate the training data; see below). Instead, the tree is inferred directly via autoregressive generation, initialized after the special inference token. During training, the model is provided with sets of unaligned sequences paired with their corresponding ground-truth phylogenetic trees, and its parameters are optimized to predict tree structures from sequence data alone (see Methods). Once trained, the model infers a phylogenetic tree when presented with previously unseen unaligned sequences together with the inference token.

## Results

### Comparing to competitors on simulated data

We evaluated the performance of BetaInfer against several standard phylogenetic inference pipelines on simulated datasets, including, UPGMA (Sokal & Michener, 1958), NJ (Saitou & Nei, 1987), IQ-TREE (Nguyen et al., 2015), and RAxML-NG (Kozlov et al., 2019). Since these methods require an MSA, we used the MAFFT algorithm (version 7.450; Katoh & Standley, 2013). To ensure that the results were not dependent on the choice of alignment software, we also tested an additional aligner (Clustal Omega; Sievers & Higgins, 2018), with the results provided in Supplementary Information S1. The protein test data included 1,000 trees with 10-14 taxa (Dataset DPT, see Supplementary Information S2). UPGMA had the lowest performance, with a normalized RF (nRF; Robinson & Foulds, 1981) distance of 0.302 ; 0.168. NJ improved substantially (with nRF of 0.088 ; 0.1), while IQ-TREE and RAxML-NG achieved even lower nRF distances of 0.062 ; 0.083. BetaInfer slightly outperformed all competitors, achieving the lowest nRF distance of 0.058 ; 0.084, demonstrating its ability to accurately reconstruct phylogenetic trees. Statistical analysis confirmed that BetaInfer’s improvement was significant compared to UPGMA and NJ (paired t-test; p ć 10^—10^), but not significant compared to IQ-TREE and RAxML-NG (paired t-test; p = 0.19 and 0.2, respectively). Overall, these results indicate that BetaInfer matches or slightly exceeds the performance of state-of-the-art phylogenetic inference methods while providing a robust, alignment-free alternative.

### Performance on out of distribution scenarios

We next evaluated the performance of BetaInfer under two out-of-distribution scenarios, in which the test data deviated from the evolutionary model encountered during training. In the first scenario, we tested the effect of substitution models. Specifically, BetaInfer was trained on simulated MSAs generated under the WAG amino-acid replacement model, the test data MSAs were generated using different replacement models, such as LG, JTT, and even matrices specific for proteins in the mitochondria or in plastids (Fig. 2). The results show that nRF distances remained relatively stable across different substitution models, indicating that the model can infer accurate tree topologies even when the underlying substitution process differs from the training distribution (Fig. 2). Statistical analysis revealed that only the MTREV24 and HIVW substitution models showed a significant difference compared to the training model (t-test; p ć 0.02), while no significant differences were observed for the other models. As expected, the three substitution models associated with the lowest performance, CPREV45, MTREV24, and HIVW are specialized models derived from specific protein classes (proteins encoded in organelles with different genetic codes than the nuclear proteins) or viral proteins. This is in contrast to the more general models such as WAG (the training model), JTT, LG, and DAY, which were estimated to reflect that amino-acid replacement rates characterizing a diverse set of nuclear-encoded MSAs.

**Figure 2:**
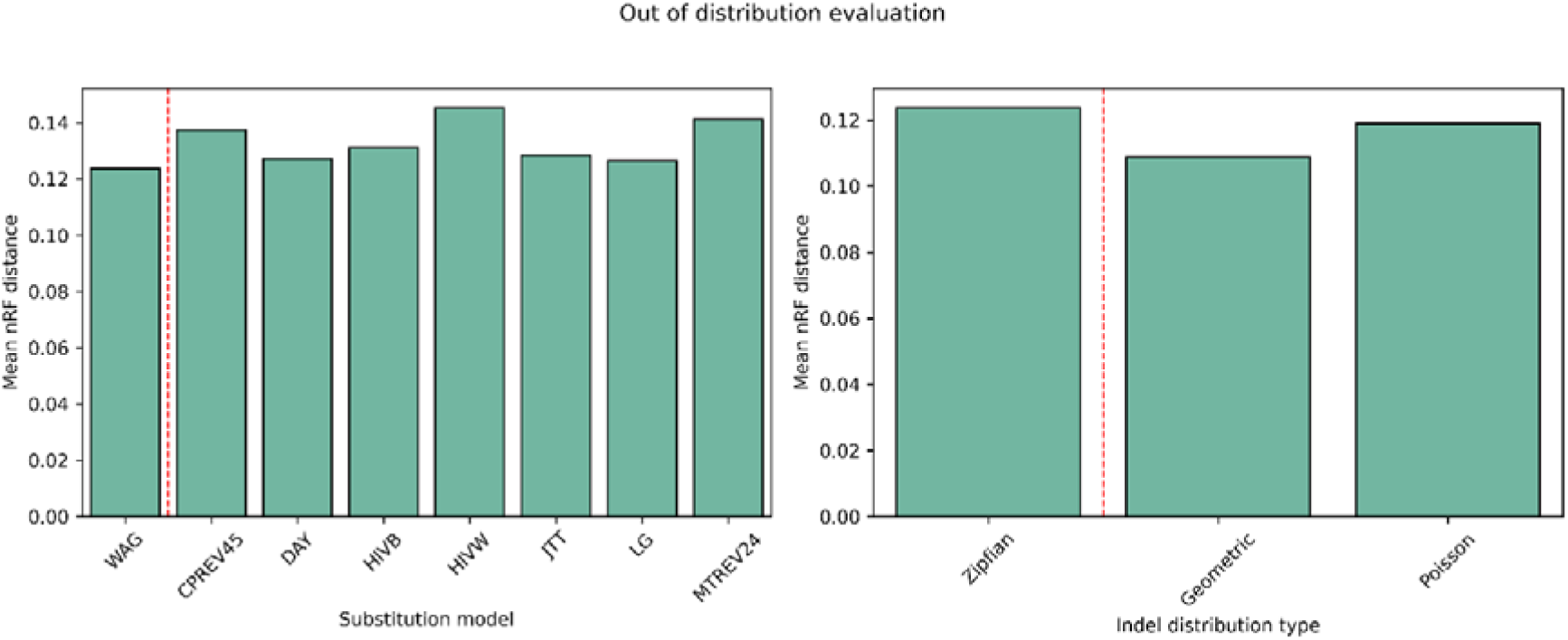
Out-of-distribution evaluation of BetaInfer. Normalized Robinson Foulds (nRF) distances for phylogenetic trees inferred from sequences evolved under substitution models or indel distributions that differ from the training distribution (see Supplementary Information S2). The dashed (red) line indicates the baseline nRF distance for the training distribution (in-distribution). Statistical analysis (t-test) showed that performance differences were significant for the MTREV24 and HIVW substitution models relative to the WAG training model and for the geometric distribution relative to the Zipfian training model; differences in all other comparisons were not statistically significant.

In the second scenario, we study the effect on the assumed indel model. BetaInfer was trained on simulated data, in which the indel size distribution was Zipfian. We assessed robustness by evaluating the performance on test data, in which the indel lengths follow a different size distribution, specifically, either geometric or Poisson. The test data included 1,000 MSAs for each indel-size distribution (see Supplementary Information S2 for a detailed description of these test datasets). Surprisingly, the performance on MSAs generated with indel models that are different than the one used for training was better compared to the performance on test data with the same distribution as the training one. This improvement was statistically significant for the geometric distribution, but not for the Poisson distribution (t-test; and , respectively). This is likely explained by the fact that the Zipfian indel distribution creates more challenging data due to longer indels. Indeed, the difficult of the inference analysis, as estimated using the approach of Haag et al. (2022) confirmed that the geometric distribution was the least difficult (0.029), while Poisson and Zipfian distributions were characterized by higher values (0.0454 and 0.0455, respectively). These results demonstrate that BetaInfer generalizes well to inference of phylogenetic trees under a variety of substitution and indel models, which is crucial for diverse datasets.

### Comparing to competitors on empirical data

To evaluate performance on empirical data, we analyzed protein families from five mammalian orders (*Artiodactyla*, *Primates*, *Chiroptera*, *Carnivora*, and *Rodentia*) using the OrthoMaM dataset, version 12 (Allio et al., 2024). For each protein family, we randomly sampled two sequences from each order, resulting in ten species per protein family. We then reconstructed a phylogenetic tree from these ten protein sequences. Accuracy was evaluated by comparing the reconstructed tree to the established mammalian species tree on 1,816 protein families that satisfied our length and coverage criteria (see Methods). Across all families, BetaInfer achieved a mean nRF distance of 0.2844, comparable to leading likelihood-based methods (RAxML-NG and IQ-TREE, both with nRF distances of 0.2837) and slightly better than NJ (RF distance of 0.2932), while outperforming UPGMA (RF distance of 0.4061). Statistical comparisons confirmed that BetaInfer’s performance was indistinguishable from RAxML-NG and IQ-TREE (paired t-test; p = 0.83), but significantly better than UPGMA and NJ (paired t-test; p ć 0.02). These results indicate that BetaInfer matches state-of-the-art phylogenetic pipelines (MSA inference followed by a heuristic maximum-likelihood search) on empirical mammalian data without relying on explicit alignment or parametric evolutionary models.

When stratifying results by total sequence length, two complementary trends became apparent. Shorter protein families posed a greater challenge for all methods, reflecting limited phylogenetic signal; in this regime, NJ performed unexpectedly well and, in some cases, even outperformed RAxML-NG and IQ-TREE (Fig. 3). As sequence length increased, likelihood-based methods showed steady improvements, whereas BetaInfer exhibited a more modest gain, suggesting that very long sequences, while informative, are more challenging to fully exploit within the current generative framework. Nonetheless, across all length ranges, BetaInfer remained competitive with established methods and consistently outperformed simple distance-based clustering (UPGMA), highlighting its robustness across heterogeneous empirical datasets.

**Figure 3:**
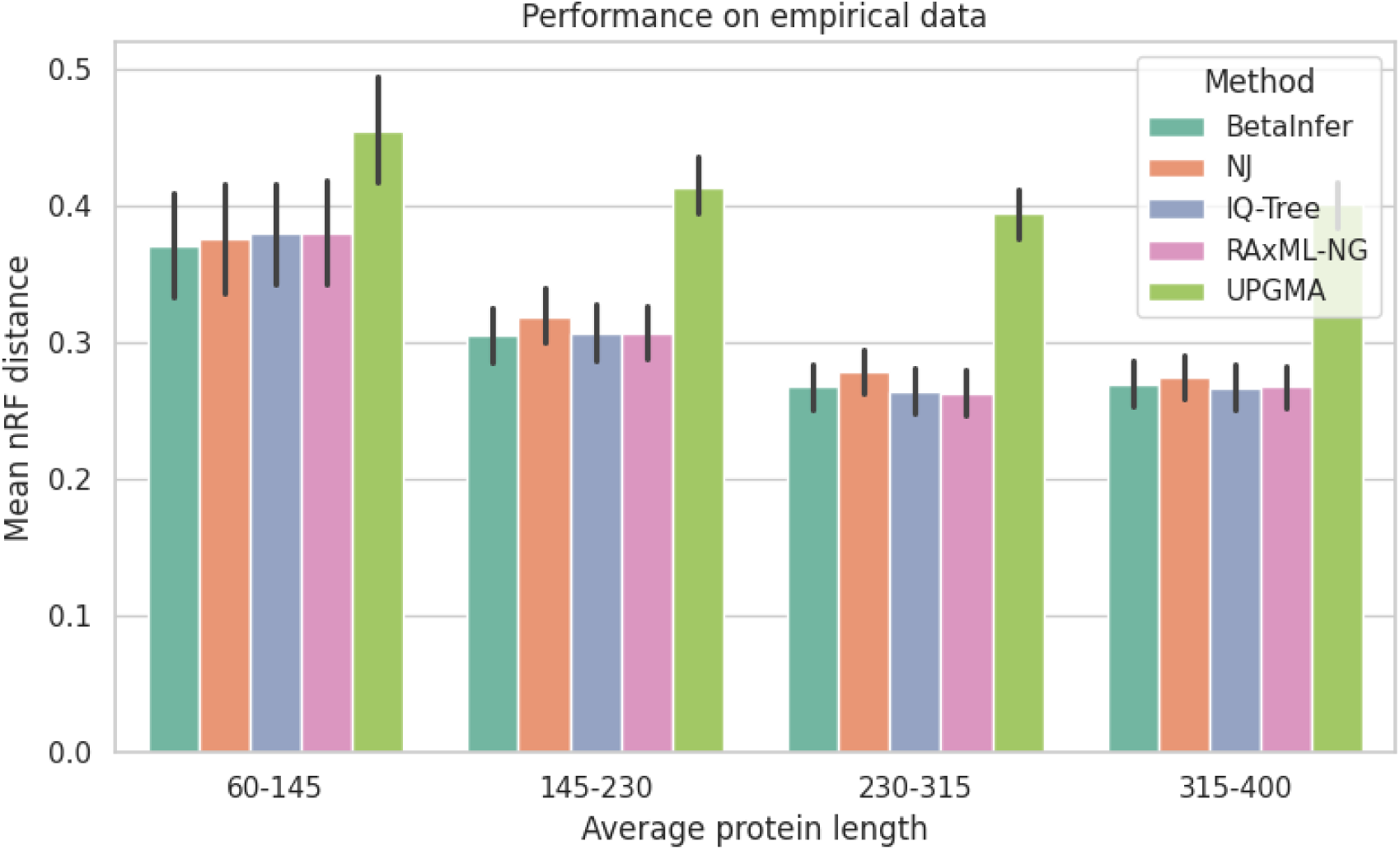
Performance of multiple phylogenetic inference methods (BetaInfer, NJ, IQ-TREE, RAxML-NG, and UPGMA) on empirical mammalian protein families, stratified by total sequence length. Trees were inferred from ten protein sequences per family. Two sequences were sampled from each of the following orders: Artiodactyla, Primates, Chiroptera, Carnivora, and Rodentia, using the OrthoMaM database (see Methods).

### Interpretating BetaInfer

One of the most difficult aspects of deep-learning algorithms is to obtain some understanding of inner working of the computational model. Through a series of hidden-layer transformations, the model refines the representation of the unaligned sequences until it reaches the output head, where it is mapped into discrete tokens, which in our case is a Newick characterizing of the evolutionary relationships. Here, we aimed to understand how the model captures the evolutionary relationships among the input sequences through the model layers. Specifically, we aimed to understand whether the model progressively learns the equivalent of distances among the input sequences. To this end, we analyzed the hidden-state representations of the model layers and evaluated whether they could be reduced to pairwise distance matrices capable of recovering the inferred trees. An evolving understanding of the evolutionary distances among the input sequences along the layers should correspond to an implicit distance-based inference within the model.

To test this hypothesis, we trained a shallow regressor to predict pairwise evolutionary distances from hidden-state representations at each layer, and applied NJ to the predicted distance matrices (see Methods). This structural probing approach have previously been used in NLP to uncover latent syntactic structures in text language model representations (Hewitt & Manning, 2019; Maudslay et al., 2020) and was recently suggested for reconstructing phylogenetic trees (Nesterenko et al., 2025; Zhang et al., 2025). Fig. 4 presents the nRF distance between the inferred tree from each layer and the true tree (Dataset DPT, see Supplementary Information S2). In the first three layers, the embeddings do not seem to encode evolutionary aspects among the sequences, as the nRF distance is close to 1. From the fourth layer onward, the nRF distance decreases, reaching a minimum of 0.2 in layer 19. The nRF distance then increases back to around 1 in the next layers. These last layers likely specialize in next-token prediction and thus emphasize local token selection over global evolutionary structure, echoing findings about stages of inference in text language models (Geva et al., 2023; Katz & Belinkov, 2023; Lad et al., 2025). While a performance gap remains between the generated trees and those derived from structural probes, these findings suggest that BetaInfer encodes sequences into a latent space and employs an implicit clustering mechanism based on learned distance metrics to reconstruct the phylogeny.

**Figure 4:**
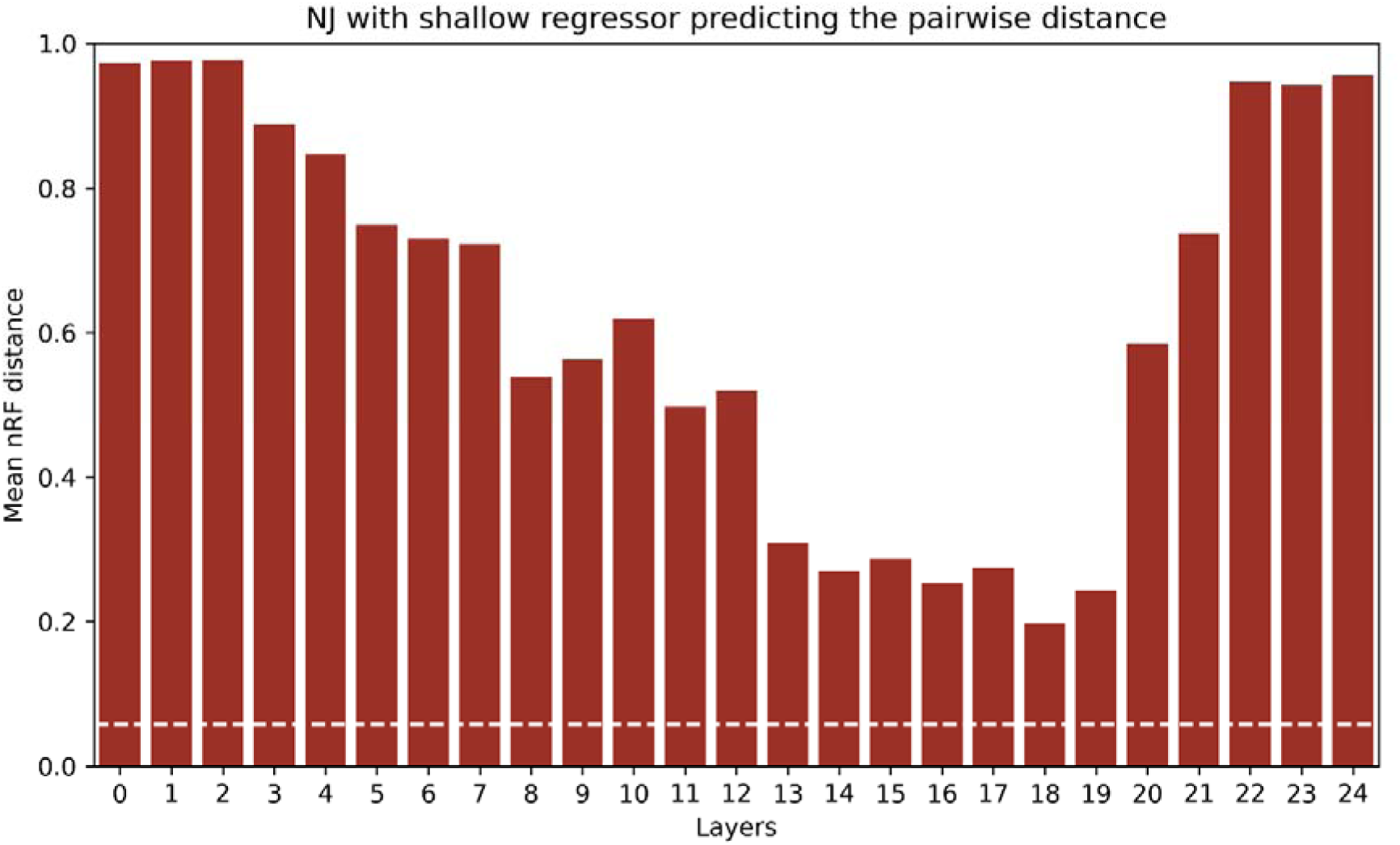
Reconstruction of phylogenetic trees from hidden-state representations using a trained pairwise-distance regressor. For each layer of our hybrid model, sequence embeddings were extracted and passed through a shallow regressor to predict pairwise distances. These predicted distances were then used by NJ to infer phylogenetic trees and assess the recoverability ofthe ‘true’ topology. The dashed (white) line ( ) indicates the nRF distance ofthe generated trees by BetaInfer.

### Optimizing performance

The above results were obtained using an optimized BetaInfer. Here we describe how we systematically evaluated several methodological choices to optimize BetaInfer performance. We began with tokenization, given that previous studies have shown it to be a critical factor for both model performance and memory efficiency (Dotan et al., 2024; Lindsey et al., 2025; Sun et al., 2025). Accordingly, we employed data-driven tokenizers (Sennrich et al., 2016) trained specifically on evolutionary sequence data and systematically assessed their impact on the number of resulting tokens. Next, we compared three model architectures, inspired by commonly used text-based language models: an SSM (Mamba2; Gu & Dao, 2024), an attention-based transformer (LLAMA; Dubey et al., 2024), and a hybrid SSM-transformer (Zamba; Glorioso et al., 2024). For each architecture, we conducted a hyperparameter search to identify the configuration that maximized performance, ultimately selecting the Zamba (hybrid) architecture. We then trained three Zamba configurations with approximately 200 million parameters and chose Configuration 1 (24 layers with a hidden size of 1,024), for further experiments. Comprehensive details of all optimization procedures are provided in Supplementary Information S3.

#### Choosing the best phylogenetic tree from multiple candidates

We evaluated a novel inference strategy in which multiple phylogenetic tree candidates are generated for the same set of unaligned proteins and the most consistent tree is subsequently selected. Specifically, if one changes the order of the unaligned sequences provided as input to BetaInfer, one may obtain a different tree. Thus, we run BetaInfer with multiple randomized orders of the sequences and obtain several candidate trees. From these trees, we select that tree with minimal nRF distance to the rest of the trees (see Methods). This approach substantially reduces reconstruction error relative to a single prediction (Table 1). Increasing the number of candidates from one to five reduces the nRF distance from 0.087 to 0.065, corresponding to a 25.2% reduction in error. Expanding the number of candidates to 15 further decreases the nRF distance to 0.061 (29.8% reduction), and with 30 candidates, the error reaches 0.059 (32.1% reduction). Beyond this point, performance gains largely saturate, with 50 candidates yielding only a marginal additional improvement (nRF distance of 0.058). Overall, these results illustrate the well-known benefit of ensemble strategies in reducing variance and improving robustness, demonstrating that aggregating multiple predictions can significantly enhance phylogenetic inference accuracy (Dotan et al., 2025; Edgar, 2022).

**Table 1:** Effect of generating multiple candidate trees and selecting the most consistent tree on phylogenetic inference accuracy. We report the mean nRF distance obtained when selecting the best tree from varying numbers of generated candidate phylogenetic trees (Dataset DPT; see Supplementary Information S2).

| Candidates<br>number | 1 | 5 | 15 | 30 | 50 |
| --- | --- | --- | --- | --- | --- |
| nRF distance | 0.087 | 0.065 | 0.061 | 0.059 | 0.058 |

### Runtime analysis

We benchmarked the computational efficiency of BetaInfer by measuring the average inference time per tree on the DPT dataset (see Supplementary Information S2) relative to several established phylogenetic pipelines. To ensure a comprehensive comparison, we measured total execution time, including all necessary preprocessing steps. UPGMA, NJ, IQ-TREE, and RAxML-NG were executed on an AMD EPYC 7H12 64-Core Processor, while BetaInfer utilized a single NVIDIA RTX A6000 GPU. UPGMA was the most efficient, requiring an average of 0.64 seconds per tree inference, followed by IQ-TREE (4.33 seconds), NJ (4.62 seconds), and RAxML-NG (28.54 seconds). BetaInfer averaged approximately four minutes (228 seconds) per inference; this duration was primarily driven by the evaluation of fifty candidate trees per unaligned protein set, which accounted for 226 seconds of the total time. Of note, unlike classic frameworks, BetaInfer requires a one-time prior training phase.

## Discussion

In this work, we introduce BetaInfer, a novel deep learning framework that reformulates phylogenetic inference as a sequence modeling problem and infers tree topologies directly from unaligned sequence data. By leveraging advanced generative modeling capabilities and an end-to-end deep learning objective, BetaInfer fundamentally departs from classical phylogenetic pipelines. Conventional methods typically rely on three core components: an MSA, an explicit model of sequence evolution (including substitution and indel), and a tree-search procedure, each of which introducing its own sources of error and computational burden. Our results demonstrate that this data-driven, generative approach can recover accurate phylogenetic trees across a wide range of simulated and empirical scenarios.

Effectively handling long biological sequences is a critical requirement for analyzing evolutionary data, particularly when working with large numbers of taxa or long protein sequences. Standard attention-based transformers are limited by the quadratic scaling of memory requirements with sequence length, making naïve application to phylogenetic data impractical. Our results highlight that both tokenization and architectural choices are essential for overcoming these constraints. In particular, data-driven tokenization plays a central role by substantially reducing the number of tokens. In addition, hybrid architectures that integrate state-space layers with attention mechanisms provide better scalability and performance. The superior performance and efficiency of such hybrid architectures demonstrate that such designs are well suited for large-scale phylogenetic inference, and suggest that future progress in this domain will directly impact biological models.

Our interpretability analysis highlights both the promise and the inherent challenges of decoding the internal mechanisms of deep learning models. We demonstrate that the intermediate hidden states of BetaInfer recover phylogenetic signals that can be mapped directly to pairwise distances. This suggests the model internalizes evolutionary relationships in a manner analogous to distance-based frameworks. However, more sophisticated interpretability techniques are needed to dissect the specific distance-based functions the architecture employs or to determine how it learns distinct substitution probabilities. Advancing these tools may reveal novel computational principles that complement or extend classical phylogenetic theory.

A foundation model in NLP is a large machine-learning model trained on vast, unlabeled text corpora usually using self-supervised learning. Rather of being optimized for a single task, it internalizes broad linguistic patterns that facilitate adaptation to many downstream applications. A critical frontier is the development of such foundation models tailored to evolutionary sequence data. Such models could learn increasingly abstract and transferable representations of evolutionary signals across diverse organisms, molecular types, and evolutionary regimes. These foundation models could serve as general-purpose engines for evolutionary analysis, processing multiple unaligned sequences to produce high-quality numeric representations.

Currently, BetaInfer is limited to few a few dozen sequences, due to the memory limitations imposed by the transformer architecture. However, emerging technologies are likely to mitigate these scaling restrictions, enabling the training of models with increased number of tokens (both in terms of sequence count and length) to meet the requirements of modern phylogenomic analyses. Notably, despite the higher inference times, BetaInfer’s deep learning architecture offers a significant advantage in theoretical algorithmic complexity, as it avoids explicit tree search, whereas traditional maximum-likelihood approaches suffer from combinatorial explosion due to the exponential increase in the tree space they must evaluate. Furthermore, BetaInfer bypasses the notoriously difficult MSA inference step. We anticipate that machine-learning generative approach such as the one presented in this study will emerge as a powerful tool for complex molecular evolution inference tasks.

In a broader sense, this study underscores the growing synergy between NLP and biological research. A key characteristic of biological data is its sequential nature: DNA, RNA, and proteins, are represented as ordered strings of symbols. NLP offers a mature and flexible set of tools for modeling such data, enabling the extraction of complex, long-range dependencies without requiring explicit heuristics. By framing core evolutionary tasks as generative problems, we demonstrate how advances in NLP can be directly translated into novel biological methodologies. This integration opens the door to unified, data-driven frameworks in which diverse biological inference problems can be addressed using advanced generative models and learning objectives. As sequencing technologies continue to expand the scale and scope of available data, the continued integration of NLP and biology is well positioned to become a central driver of progress, reshaping how evolutionary relationships and molecular processes are inferred from sequence data.

## Supporting information

Supplementary Information: Phylogenetic tree inference using generative models

## Data availability

Code and models are available on our GitHub page: https://github.com/technion-cs-nlp/BetaInfer.

## Materials and methods

### Outline

We begin by describing in detail how deep learning can generate phylogenetic trees. We then extend this framework to ancestral sequence reconstruction (ASR) and MSA inference, which are integrated into the training procedure. Next, we address the challenges posed by long biological sequences and present solutions based on tokenization strategies and hybrid model architectures. We subsequently describe the training protocols and datasets used throughout the study. We then introduce our approach to improving inference performance by generating and selecting among multiple candidate phylogenetic trees. Finally, we discuss the evaluation metrics used to assess model performance across all experiments.

### Generative approach for phylogenetic tree inference

A wide range of deep learning architectures can be used for sequence generation; among them, decoder-only transformer models are particularly effective, achieving strong performance through autoregressive next-token prediction (Radford et al., 2018). In this work, we leverage such models to generate phylogenetic trees (formatted via the Newick format). Tree generation is framed as a series of multiclass prediction steps, in which the model predicts one token at a time, corresponding to elements of a linearized tree representation (e.g., topology and branch-related tokens). The input sentence (i.e., unaligned proteins) is first tokenized into substrings represented as integers, which are then mapped into a high-dimensional embedding space via an embedding layer. These embeddings are processed by a stack of layers, producing at the last layer a probability distribution over the vocabulary via a Softmax function. A vocabulary item (i.e., token) is selected from this distribution according to a specified decoding strategy (e.g., greedy decoding), and is appended to the input sequence. This autoregressive process continues until a predefined termination criterion is met, or until the generation of a special end-of-sequence token (“<EOS>”).

### ASR and MSA as a next token prediction task

Both ASR and MSA inference can be naturally reformulated as next-token prediction problems by casting evolutionary inference as a sequence completion task. As in phylogenetic tree inference, the unaligned sequences are first converted into a single input “sentence” using the “concat” representation (Dotan et al., 2023), in which individual sequences are concatenated by a dedicated delimiter (the pipe symbol; “|”). For ASR inference, the “<RECONSTRUCT>” token is appended to the end of this input to signal that the model should generate the ancestral sequence. For MSA, the target output is an aligned sequence representation. To enable autoregressive generation of an alignment, the MSA is linearized into a single sequence using the “spaces” format, which encodes the alignment column by column: for an alignment of N sequences, the first N amino-acids correspond to the first alignment column, the next N amino-acids to the second column, and so on (Dotan et al., 2025). A dedicated token, “<ALIGN>”, is appended after the concatenated input to specify that the task is alignment. Under this formulation, phylogenetic tree inference, ASR and MSA reduce to predicting a sequence of tokens conditioned on the input context, allowing the same transformer architecture and training objective to be applied across tasks by modifying the task-specific control token and output representation. We recently explored the use of generative models for ASR in a separate preprint (Dotan et al., 2026).

### Handling long sequences

Inferring phylogenetic trees directly from unaligned biological sequences poses substantial computational challenges due to both the length and number of sequences involved. Attention-based transformer architectures scale quadratically in memory and computation with respect to the number of tokens, which typically limits practical training and inference to a few thousand tokens (Lin et al., 2021). In the context of phylogenetic tree inference, the input often consists of dozens of protein sequences, each potentially spanning hundreds or thousands of residues, yielding input lengths that typically exceed the capacity of the attention-based models. To address these limitations, we adopt a combination of architectural and representational strategies that enable efficient modeling of long sequence inputs while preserving inference accuracy.

Specifically, we employ hybrid architectures that integrate SSM components with attention-based layers. SSMs have recently emerged as a scalable alternative to full attention, as their layers summarize historical information using a compact state representation whose memory requirements scale linearly with input length, rather than quadratically (Gu et al., 2022; Gu & Dao, 2024). While pure SSM-based models offer substantial efficiency gains, attention mechanisms are generally more effective at capturing complex, long-range dependencies that are critical for accurate evolutionary inference (Nguyen et al., 2023). Hybrid architectures, such as Zamba (Glorioso et al., 2024) and Jamba (Lenz et al., 2024), combine these complementary strengths by interleaving SSM blocks with attention layers. This design allows the model to efficiently process longer sequences while retaining the expressive power of attention for modeling complex patterns, making hybrid models particularly well suited for large-scale phylogenetic tree inference.

In addition to architectural choices, we reduce the effective sequence length through data-driven tokenization. While many biological language models represent sequences using single-amino-acid tokens, we leverage multi-amino-acid tokenization to improve both computational efficiency and predictive performance (Dotan et al., 2024). Tokens correspond to characters or substrings that map discrete input sequences to high-dimensional embeddings processed by the model and later decoded back into symbolic form (via the Softmax function). We trained multiple Byte-Pair Encoding (BPE) tokenizers (Sennrich et al., 2016) with vocabulary sizes ranging from 100 to 25,600, using a subset of the training data (DT; see Supplementary Information S2). The BPE algorithm iteratively merges frequently co-occurring token pairs, producing a data-driven vocabulary in which individual tokens may represent short motifs or multi-residue segments. For phylogenetic tree inference, BPE was applied to the unaligned sequence inputs, while the tree outputs, represented in Newick format, were encoded using single-character tokens, e.g., “(”, “)”, “,”, “:”, etc. By learning compact representations of recurring sequence patterns, this tokenization scheme substantially reduces input length by more than a factor of two, enabling efficient processing of long and numerous biological sequences.

### Training

We adopted a training strategy that relies on simulated data to learn phylogenetic tree inference. Simulations provide access to ground-truth labels, including true tree topologies and branch lengths, enabling direct supervision that is not available for empirical datasets. In this setting, the model was trained to predict phylogenetic trees from unaligned sequences using labels generated by the simulations (see below). Model parameters were optimized by minimizing the cross-entropy loss over the tree representations, allowing the model to directly learn the mapping from protein sequences to underlying evolutionary structure. Training and evaluation were implemented using the HuggingFace framework (Wolf et al., 2020). Specific training parameters are provided in Supplementary Information S4.

#### Transfer learning

Our training framework makes extensive use of transfer learning, enabling knowledge acquired from simpler inference settings to facilitate learning in more challenging ones (Avram et al., 2025; Kim et al., 2022; Zhuang et al., 2021). Specifically, model parameters optimized on an easier dataset were used to initialize training on progressively harder datasets. In the first phase, dataset difficulty was controlled through simulation parameters, primarily the branch length, with longer branches corresponding to more difficult phylogenetic reconstructions. Training began with simulations capped at 0.05 substitutions per site and the maximum branch length was increased linearly to 0.2 substitutions per site. A detailed description of the datasets is provided below.

### Data

#### Simulated data

To train and evaluate BetaInfer, we generated large synthetic datasets containing unaligned sequences along with their corresponding true alignments, ancestral sequences, and phylogenetic trees. The datasets were produced using SpartaABC (Loewenthal et al., 2021; Wygoda et al., 2025), a flexible simulator that allows precise control over evolutionary parameters. For each dataset, sequences evolved along randomly generated trees with diverse branch lengths and topologies. The simulator models indels asymmetrically, in both rate and length distribution, providing a realistic representation of molecular evolutionary complexity. These datasets enabled consistent training and evaluation of model performance. A detailed description of the simulations and parameter settings is provided in Supplementary Information S2.

#### Empirical data

To evaluate performance on empirical data and compare different phylogenetic inference pipelines, we used the OrthoMaM dataset, version 12 (Allio et al., 2024), which contains orthologous protein families from mammalian species. We focused on five well-sampled mammalian clades, *Artiodactyla*, *Primates*, *Chiroptera*, *Carnivora*, and *Rodentia*. We considered the 6,555 protein families that contain at least two sequences from each of the five clades. For each protein family, we randomly sampled two protein sequences from each clade, yielding 10 unaligned sequences per protein family. Because sequence length poses computational challenges (for BetaInfer), we further restricted the analysis to families for which the total number of amino acids across all sampled sequences was less than 4,000, resulting in 1,816 protein families retained for downstream analyses. We compared the inferred phylogenetic trees to the accepted mammalian species tree, which is well established (Arnason et al., 2002; Murphy et al., 2001), providing a reference for topological accuracy. Specifically, the inferred trees were compared to the following reference topology: “((*Primates*, *Rodentia*), (*Chiroptera*, (*Carnivora*, *Artiodactyla*)));”.

### Generating multiple candidates and choosing the best phylogenetic tree

It is possible to infer multiple phylogenetic trees from the same set of sequences, increasing confidence in results and reducing sensitivity to modeling choices. For example, alternative phylogenies can be generated by varying guide trees, model parameters, or heuristic search strategies, and such ensembles are commonly used to assess topological robustness and branch support (e.g., bootstrap or Bayesian posterior sampling). Similar techniques have been tested and successfully implemented in the MSA inference (Dotan et al., 2025; Edgar, 2022). Inspired by these approaches, we developed an analogous strategy for generating multiple phylogenetic trees within the deep learning framework proposed here.

Specifically, our approach exploits the fact that the model operates on a concatenated representation of unaligned input sequences. By permuting the order of sequences provided to this “concat” representation, we induce variability in the inferred evolutionary relationships (Dotan et al., 2025), resulting in alternative phylogenetic trees for the same input set. For example, a dataset containing three sequences admits six possible input permutations, each of which can lead to a distinct inferred tree topology. More generally, for larger datasets, this procedure yields a diverse collection of alternative phylogenetic reconstructions.

Formally, let S, T, and f denote a list of unaligned sequences, a phylogenetic tree and a phylogenetic inference procedure, respectively. The phylogenetic tree procedure depends on a set of model parameters or configurations, denoted by θ. Varying θ produces different phylogenetic trees for the same input S. Given a collection of k configurations: θ_1_, θ_2_, … , θ_k_, the procedure yields a corresponding set of inferred trees: f_θ1_ (S), f_θ2_ (S), … , f_θk_ (S) which we denote as T_1_ , T_2_ , … , T_k_. Of note, some configurations may converge to identical trees. In our framework, alternative configurations are obtained by permuting the order of the unaligned input sequences. For each inferred tree T_f_ , we compute a tree certainty score by comparing it to all other trees in the set, based on the average agreement, measured by nRF. The chosen tree is the one that maximizes this certainty measure, i.e., the phylogenetic tree that minimize the nRF distance to all other trees.

### Comparison to competing methods

We evaluated several phylogenetic inference methods: UPGMA (Sokal & Michener, 1958), NJ (Saitou & Nei, 1987), IQ-TREE (version 2.2.2.6; Nguyen et al., 2015), and RAxML-NG (version 1.2.0; Kozlov et al., 2019). Each pipeline was tested using two sequence alignment tools: MAFFT (version 7.450; Katoh & Standley, 2013) and Clustal Omega (version 1.2.4; Sievers & Higgins, 2018).

### Probing hidden-state representations for reconstructing trees with pairwise distance

To assess whether phylogenetic relationships can be recovered from the internal representations of BetaInfer, we conducted a probing analysis in which simple supervised models were trained on internal representations to test whether specific evolutionary information is encoded in these representations. This approach is inspired by representational probing methods widely used in NLP (Adi et al., 2016, p. 201; Belinkov, 2022; Belinkov & Glass, 2019; Hupkes et al., 2020; Pimentel et al., 2020). For each model layer, we extracted the hidden-state embeddings corresponding to each unaligned input sequence. Dataset DPT (see Supplementary Information S2) was used for this analysis. Each sequence was represented by a fixed-length vector, obtained by average pooling across its token embeddings. Using these representations, we evaluated a strategy for deriving pairwise sequence distances by training a lightweight supervised regressor to map hidden-state vector pairs to evolutionary distances. This regressor comprising a 2,048-neurons layer with ReLU activation, followed by a 256-neurons output layer. It was trained using labels derived from ground-truth phylogenetic trees to map hidden states to pairwise distances. The (regressor) dataset comprised 60,000 pairs, each consisting of fixed-size vectors representing two unaligned sequences along with the corresponding phylogenetic distance between them. The data were split into 50,000 and 10,000 pairs for training and validation, respectively. The regressor was trained independently for each layer, thereby probing the extent to which phylogenetic distance information is linearly or nonlinearly accessible at different depths of the model. The resulting distance matrices were then used as input to the NJ algorithm to reconstruct phylogenetic trees, which were compared to the true trees using nRF distance. This probing framework allowed us to systematically evaluate whether, and at which layers, phylogenetic information is encoded in a form compatible with distance-based tree reconstruction.

### Cross-entropy loss

The model generates a token by calculating a probability distribution over the full tokenizer vocabulary. Cross-entropy loss quantifies the difference between this predicted distribution and the true target distribution, in which the correct token has probability one and all other tokens have probability zero. The loss for a given token is defined as the negative logarithm of the probability assigned to the correct token. The loss for a sequence is computed by averaging this value over all predicted positions, and the dataset-level loss is obtained by further averaging across all sequences. Lower cross-entropy values indicate that the model assigns higher probability to the correct tokens, achieving more accurate predictions. Training loss is used to optimize the model weights and validation loss is used to monitor performance, choose best hyperparameters and detect overfitting.

### Tree inference evaluation

The RF distance (Robinson & Foulds, 1981) quantifies the dissimilarity between two phylogenetic trees by counting the number of partitions that differ between them. We employed this metric to benchmark the various approaches, comparing inferred trees against both simulation-based ground truths and established mammalian taxonomies. Lower RF distances indicate better topological agreement, thus reflecting improved evolutionary inference performance. We normalized the RF (nRF) distance by dividing the score with the total number of splits, thus the range of this metric is between zero and one. Of note, since some inference approaches return unrooted trees, we report the nRF distance computed on unrooted trees (if an approach returns a rooted tree, it is transformed to unrooted before comparison).

In addition to topological accuracy, we assessed whether the model produced structurally valid phylogenetic trees in Newick format. A valid Newick tree must satisfy several constraints: (1) parentheses must be properly balanced to define hierarchical clades; (2) commas must correctly separate sibling subtrees within the same clade; (3) colons must appear only after node labels to denote branch lengths; (4) and each leaf must appear exactly once, ensuring a one-to-one correspondence between input sequences and leaf nodes.

Violations of these rules result in invalid trees that cannot be meaningfully compared to a reference topology. In the inference stage, we obtain multiple candidate trees (generated by permuting the sequence input order, see above). If an output Newick format is invalid, we assign a maximal nRF distance of 1.0 between this tree and all other candidate trees. This procedure ensures that this tree will never be selected among the candidate trees.

