## Supplementary Information: Phylogenetic tree inference using generative models for "Phylogenetic tree inference using generative models"

### Supplementary Information S1: Running the competitors with an additional aligner

We repeated the comparison of phylogenetic inference methods using sequences aligned with Clustal Omega (Sievers & Higgins, 2018) to verify that our results were not dependent on the choice of alignment software. Overall trends were similar to those observed with MAFFT, with a slight decrease in performance for all MSA-based methods. UPGMA, NJ, IQ-TREE, and RAxML-NG had an nRF distance of 0.303, 0.091, 0.064, and 0.063, respectively (Table S6). BetaInfer’s performance remained unchanged, as our approach does not depend on an alignment input. Statistical analysis indicated that the differences between BetaInfer and UPGMA, NJ, and IQ-TREE were significant (paired t-test, $p < 0.032$), while the difference compared to RAxML-NG was not significant ($p = 0.05003$).

*Table S6: Comparison of phylogenetic reconstruction accuracy on simulated data when using the Clustal Omega aligner. We report the performance of: UPGMA, NJ, IQ-TREE, RAxML-NG, and BetaInfer on Dataset DPT (see Supplementary Information S2).*

| Approach | UPGMA | NJ | IQ-TREE | RAxML-NG | BetaInfer |
| --- | --- | --- | --- | --- | --- |
| nRF distance | 0.303 | 0.091 | 0.064 | 0.063 | 0.058 |

### Supplementary Information S2: Detailed simulation description and datasets parameters

For each simulation instance, we first sampled a random phylogenetic tree containing between 10 and 14 taxa using the ETE3 library with default settings (Huerta-Cepas et al., 2016). Insertion and deletion parameters were then drawn independently to allow for a diverse indel process, with $R_{i}, R_{d}\in\left( 0.0, 0.05 \right)$ and $A_{i}, A_{d}\in(1.01, 2.0)$, where, $R_{i}, R_{d}, A_{i},$ and $A_{d}$ correspond to the insertion and deletion rates and the Zipfian length parameters for insertion and deletion, respectively (Loewenthal et al., 2021). Sequence evolution along the tree was modeled using the WAG+Γ substitution model, with a gamma shape parameter of 1.0 and four discrete rate categories. Each simulation produced one of the three types of ground-truth data: (i) the true phylogenetic tree (including topology and branch lengths); (ii) a set of unaligned sequences (including ancestral sequences at all internal nodes); and (iii) the corresponding true MSA.

Using this procedure, we first generated Dataset D1, comprising 1,080,000 simulated instances, of which 1,050,000 were used for training and 30,000 for validation. In D1, branch lengths were short (uniformly sampled in the range 0 to 0.05 substitutions per site), yielding relatively easy phylogenetic inference scenarios (Table S5). We constructed input-label pairs $(X, Y)$, where $X$ denotes the set of unaligned sequences and $Y$ denotes the target output. To support the multiple evolutionary tasks, labels were evenly distributed across three tasks: phylogenetic tree inference (encoded with the Newick format), ancestral sequence prediction at the root, and MSA prediction.

We next generated Dataset D2 by simulating an additional 1,080,000 instances with longer branch lengths, increasing the difficulty of phylogenetic reconstruction (Table S5). The model trained on D2 was initialized with the optimized weights obtained on D1, and this training strategy was iteratively extended to datasets D3 through D7, each characterized by progressively longer branch lengths and increased evolutionary divergence. We then fine-tuned the model for the phylogenetic tree prediction by simulating a DPT (“dataset phylogenetic tree”) containing only trees as labels for the unaligned sequences. Finally, for the empirical analysis, we simulated and fine-tuned BetaInfer on DSO (“dataset simulated OrthoMaM”) containing only trees as labels, but with longer protein lengths.

In addition, we constructed a separate dataset, DT (“dataset tokenizer”), for training and evaluating the tokenizer. Unlike Datasets D1 to D7, DT sampled root sequence lengths from a normal distribution with a mean of 345 amino acids and a standard deviation of 76, based on empirical protein length distributions (Nevers et al., 2023). DT comprised 7,000 instances, split into 6,000 for training and 1,000 for validation.

We also generated a dedicated out-of-distribution evaluation dataset containing 9,000 validation instances without any training data. This dataset includes 7,000 instances split equally between the following substitutions matrices: (1) CPREV45 (Adachi et al., 2000); (2) DAY (Dayhoff, 1978); (3) HIVB (Nickle et al., 2007); (4) HIVW (Nickle et al., 2007); (5) JTT (Jones et al., 1992); (6) LG (Le & Gascuel, 2008); and (7) MTREV24 (Adachi & Hasegawa, 1996), and 2,000 instances split equally between the following indel distributions: (1) Poisson $\lambda\in(0.04,0.66)$; and (2) Geometric $p\in(1.5, 25)$.

*Table S5: Simulated datasets and their parameters. We provide the normal distribution parameters for the root length: mean (*$\mu$*) and standard deviation (*$\sigma$*).*

| Datasets | Branch lengths | Number of species | Root length |
| --- | --- | --- | --- |
| D1 | (0, 0.05) | [10, 14] | $\mu=150, \sigma=30$ |
| D2 | (0, 0.075) | [10, 14] | $\mu=150, \sigma=30$ |
| D3 | (0, 0.10) | [10, 14] | $\mu=150, \sigma=30$ |
| D4 | (0, 0.125) | [10, 14] | $\mu=150,\sigma=30$ |
| D5 | (0, 0.15) | [10, 14] | $\mu=150, \sigma=30$ |
| D6 | (0, 0.175) | [10, 14] | $\mu=150, \sigma=30$ |
| D7 | (0, 0.20) | [10, 14] | $\mu=150, \sigma=30$ |
| DPT | (0, 0.20) | [10, 14] | $\mu=150, \sigma=30$ |
| DSO | (0, 0.20) | 10 | $\mu=345, \sigma=76$ |
| DT | (0, 0.20) | 14 | $\mu=345, \sigma=76$ |

### Supplementary Information S3: Optimizing performance

#### Effect of tokenizer size on the average number of tokens

In generative models for phylogenetic tree inference, tokenization converts unaligned biological sequences into sequences of discrete tokens that are mapped to high-dimensional representations processed by the model. Each token may correspond to a single amino acid or to a short k-mer spanning multiple adjacent residues (Dotan et al., 2024). Increasing the tokenizer vocabulary size allows tokens to represent longer sequence fragments, thereby reduces the total number of tokens per sequence and consequently lowers memory usage and computational cost. However, excessively large vocabularies may negatively impact inference accuracy, as the model must predict the next token from a broader and more sparse distribution. To balance these trade-offs, we employed a BPE tokenizer (Sennrich et al., 2016), which is trained to automatically determines an optimal set of tokens under a predefined vocabulary size constraint (see Methods). We evaluated BPE tokenizers with vocabulary sizes ranging from 100 to 25,600 using simulated phylogenetic datasets (Dataset DT; see Supplementary Information S2). As shown in Fig. S1, the average number of tokens per sequence decreases with increasing vocabulary size and eventually plateaus beyond a certain point. Based on this saturation behavior, we selected a vocabulary size of 6,400 for all subsequent phylogenetic inference experiments. Of note, our tokenizers are trained on unaligned input sequences, and each symbol of the Newick format is considered a separate token (see Methods).

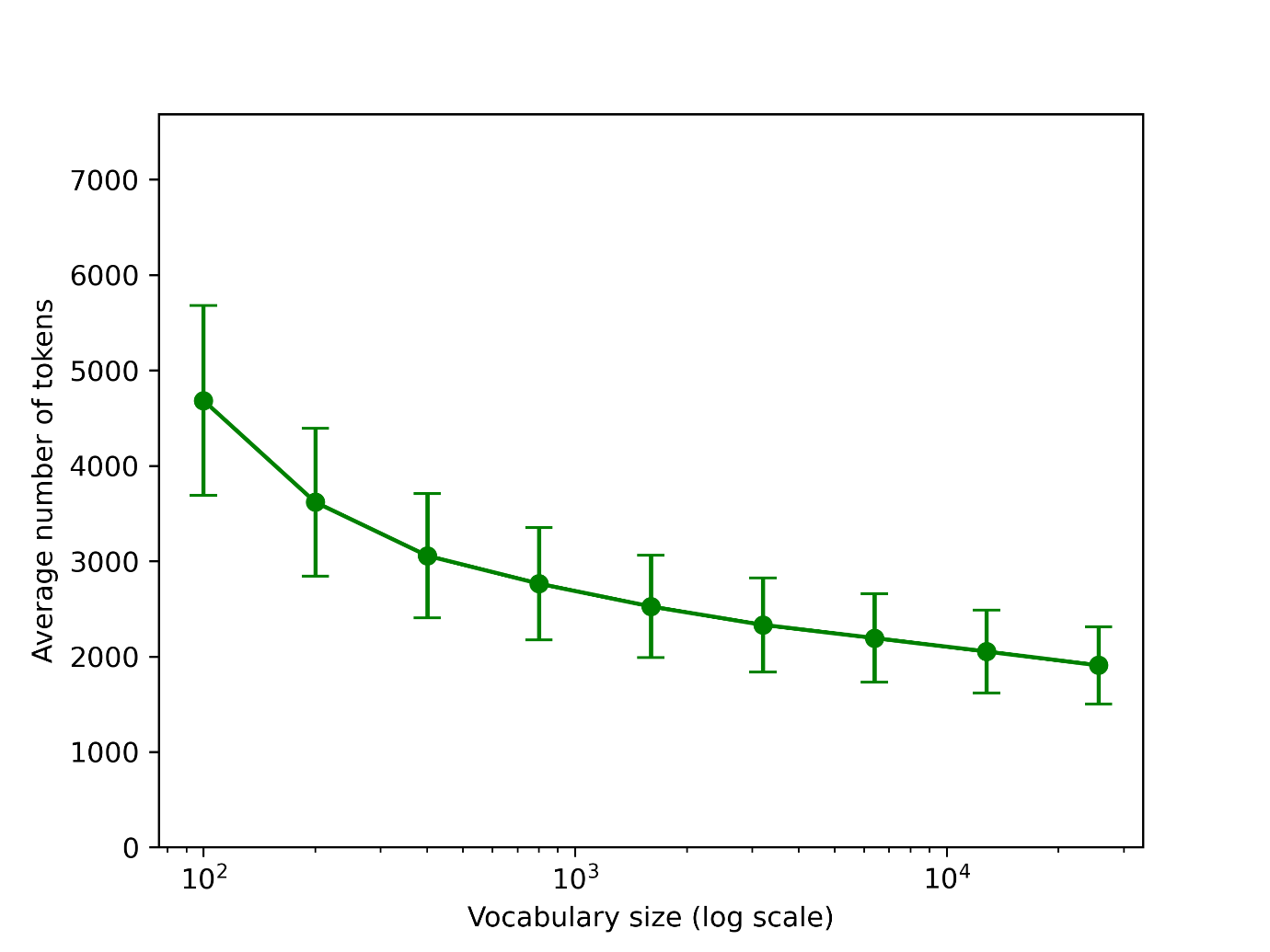

Figure S1: Effect of tokenizer vocabulary size on the average number of tokens for phylogenetic tree inference. The number of unaligned sequences is 14, with an average protein length of 345 (Dataset DT; see Supplementary Information S2).

#### Comparing the performance of different architectures

We trained and evaluated three different transformer architectures: Zamba (Glorioso et al., 2024), LLAMA (Dubey et al., 2024) and Mamba2 (Dao & Gu, 2024), each with three distinct configurations. These architectures represent complementary modeling paradigms: Mamba2 implements a state-space modeling approach, LLAMA relies exclusively on attention mechanisms, and Zamba adopts a hybrid architecture that integrates both state-space and attention layers (see Methods). To ensure a fair comparison, architecture-specific hyperparameters were optimized independently for each model (see below). We assessed model performance using three criteria: (i) validation loss, measured as cross-entropy loss on a validation set; (ii) the fraction of generated outputs that formed valid phylogenetic trees; and (iii) the nRF distance (Robinson & Foulds, 1981), computed only for trees in valid Newick format (see Methods). Although nRF distance is a standard metric for evaluating tree reconstruction accuracy when the ground-truth topology is known, its applicability at this stage was limited because the models failed to reliably generate valid early in training. This limitation is likely attributable to the relatively short training schedule (5,120 optimization steps on Dataset D1; see Supplementary Information S2). Among the evaluated architectures, Mamba2 performed poorly, exhibiting a high validation loss and failing to generate any valid phylogenetic trees (Table S1). In contrast, LLAMA and Zamba achieved comparable validation losses (1.248 and 1.405, respectively), but Zamba produced more than twice as many valid trees (200 vs. 83). Given this clear advantage, together with its more favorable memory footprint, we selected Zamba as the backbone architecture for BetaInfer.

*Table S1: Performance of different architectures in terms of validation loss, number of valid trees, and average nRF distance (computed on valid trees only). All three architectures contain approximately 200 million parameters. The results were obtained after hyperparameter optimization.*

| **Architecture** | Zamba | LLAMA | Mamba2 |
| --- | --- | --- | --- |
| **Valid loss** | 1.248 | 1.405 | 6.656 |
| **Number of valid phylogenetic trees** | 200 / 1,000 | 83 / 1,000 | 0 / 1,000 |
| **Average nRF (on valid trees)** | 0.87 | 0.85 | - |

#### Hyperparameters optimization

Table S2 summarizes the hyperparameter optimization results across model architectures (Zamba, LLAMA, and Mamba2), configurations (“deep and narrow”, “balanced”, and “shallow and wide”), and learning rates ($1\times{10}^{-4}$, and $1\times{10}^{-5})$. Overall, clear and consistent performance differences emerged among the tested architectures. Zamba systematically outperformed both LLAMA and Mamba2 across all configurations, achieving lower validation loss and producing substantially more valid phylogenetic trees. Among the three configurations, the balanced setting (Configuration 2) yielded the best results for Zamba, with the lowest validation loss (1.248 at a learning rate of $1\times{10}^{-4}$) and the highest number of valid trees (200 out of 1,000). LLAMA achieved moderate validation loss values comparable to Zamba in some settings but generated far fewer valid trees, indicating difficulties in producing structurally correct Newick outputs. In contrast, Mamba2 consistently exhibited high validation loss and failed to generate valid phylogenetic trees across all configurations and learning rates. Lowering the learning rate to $1\times{10}^{-5}$ degraded performance for all architectures, reflected by higher validation loss and a sharp drop in the number of valid trees. Together, these results underscore the importance of architectural design and hyperparameter tuning, and motivated our choice of Zamba as the backbone architecture for BetaInfer.

*Table S2: Performance comparison of model architectures, configurations, and learning rates. Models were evaluated using three performance metrics: validation loss, the number of valid phylogenetic trees generated by the model, and the average nRF distance. Each of the 18 models comprised approximately 200 million parameters and was trained on the same Dataset D1 for 5,120 training steps.*

|  | **Configuration 1 (deep and narrow)** | | | **Configuration 2 (balanced)** | | | **Configuration 3 (shallow and wide)** | | |
| --- | --- | --- | --- | --- | --- | --- | --- | --- | --- |
| **Architecture** | Zamba | LLAMA | Mamba2 | Zamba | LLAMA | Mamba2 | Zamba | LLAMA | Mamba2 |
| **Layers** | 24 | 24 | 24 | 16 | 16 | 16 | 12 | 12 | 12 |
| **Hidden size** | 1,024 | 832 | 1,024 | 1,280 | 1,024 | 1,280 | 1,408 | 1,152 | 1,536 |
| **Valid loss**  **(learning rate of** $\boldsymbol{1\times10⁻⁴}$**)** | 1.367 | 1.535 | 8.296 | 1.248 | 1.405 | 7.661 | 1.419 | 1.436 | 7.896 |
| **Number of valid phylogenetic trees**  **(learning rate of** $\boldsymbol{1\times10⁻⁴}$**)** | 102 / 1,000 | 0 / 1,000 | 0 / 1,000 | 200 / 1,000 | 71 / 1,000 | 0 / 1,000 | 117 / 1,000 | 83 / 1,000 | 0 / 1,000 |
| **Average nRF**  **(learning rate of** $\boldsymbol{1\times10⁻⁴}$**)** | 0.86 | - | - | 0.87 | 0.85 | - | 0.85 | 0.86 | - |
| **Valid loss**  **(learning rate** $\boldsymbol{1\times1}\boldsymbol{0}^{\boldsymbol{-5}}$**)** | 2.383 | 5.719 | 6.793 | 2.237 | 5.631 | 6.656 | 2.417 | 5.567 | 7.313 |
| **Number of valid phylogenetic trees**  **(learning rate** $\boldsymbol{1\times1}\boldsymbol{0}^{\boldsymbol{-5}}$**)** | 1 / 1,000 | 0 / 1,000 | 0 / 1,000 | 54 / 1,000 | 0 / 1,000 | 0 / 1,000 | 81 / 1,000 | 0 / 1,000 | 0 / 1,000 |
| **Average nRF**  **(learning rate of** $\boldsymbol{1\times1}\boldsymbol{0}^{\boldsymbol{-5}}$**)** | 0.89 | - | - | 0.82 | - | - | 0.83 | - | - |

#### Comparing different Zamba configurations

We next compared different Zamba configurations to assess the impact of model depth and width on performance. Unlike the hyperparameter experiment, models were trained longer on evolutionary data, resulting in a substantially higher proportion of valid phylogenetic trees and lower nRF distances across all configurations (Table S3). Despite comparable validation loss, clear differences emerged in topological accuracy (Table S3). Configuration 1 (deep and narrow) achieved the lowest average nRF distance (0.087), outperforming both Configuration 2 (balanced; nRF distance of 0.095) and Configuration 3 (shallow and wide; nRF distance of 0.125). While all configurations generated nearly all outputs in valid format, the improved performance of Configuration 1 indicates that increased depth is beneficial for capturing phylogenetic structure. Based on its superior nRF performance, we selected Configuration 1 for further analyses.

*Table S3: Comparison of Zamba architectural configurations for phylogenetic tree inference. The table reports model depth (number of layers), hidden dimension size, validation loss, the number of valid phylogenetic trees generated, and the average nRF distance. We report the results on Dataset DPT (see Supplementary Information S2).*

|  | **Configuration 1**  **(deep and narrow)** | **Configuration 2 (balanced)** | **Configuration 3 (shallow and wide)** |
| --- | --- | --- | --- |
| **Layers** | 24 | 16 | 12 |
| **Hidden size** | 1,024 | 1,280 | 1,408 |
| **Valid loss** | 2.406 | 2.409 | 2.416 |
| **Number of valid phylogenetic trees** | 997 / 1,000 | 999 / 1,000 | 1,000 / 1,000 |
| **Average nRF** | 0.087 | 0.095 | 0.125 |

#### Training paradigms

To assess the effect of training strategies and transfer learning on phylogenetic tree inference, we compared three training paradigms using the best-performing model (Configuration 1). In the “multiple training” strategy, a single model was trained sequentially across related evolutionary tasks (phylogenetic tree inference, MSA, and ASR), progressing from simpler datasets to more complex ones (see Methods). In contrast, the “specific training” strategy involved training a dedicated model solely for phylogenetic tree inference, using datasets of increasing complexity but without exposure to other tasks. Finally, in the “without transfer learning” setting, the model was trained from scratch directly on the most complex phylogenetic dataset.

As summarized in Table S4, models that leveraged transfer learning substantially outperformed those trained without it. “Multiple training” achieved the lowest validation loss (2.406) and the best topological accuracy, as reflected by the lowest average nRF distance (0.087). While “specific training” yielded a comparable number of valid trees and a similar validation loss, its nRF distance was higher (0.099), indicating reduced topological accuracy. In contrast, training without transfer learning resulted in a marked performance degradation, with a substantially higher nRF distance (0.513) and fewer valid trees. Notably, because the training hyperparameters and the phylogenetic dataset were fixed, the “specific training” strategy involved more training iterations on the same data than the “multiple training” strategy. Taken together, these results demonstrate that transfer learning, particularly when combined with joint training across complementary evolutionary tasks, significantly improves the accuracy of phylogenetic tree inference.

*Table S4: Effect of training strategy on phylogenetic tree inference performance. We compare three training strategies: (1) “multiple training”, which refers to a single model trained progressively on simpler to more complex datasets across all three tasks (ASR, MSA, and phylogenetic tree inference); (2) “specific training”, which denotes a model trained exclusively for phylogenetic tree inference, using datasets of increasing complexity; (3) “without transfer learning”, which corresponds to a model trained from scratch on the most complex phylogenetic dataset without prior exposure to simpler data or related tasks. All experiments were conducted using Configuration 1 (24 layers, hidden size 1,024), and performance was evaluated using validation loss, the number of valid phylogenetic trees generated, and average nRF distance. We report results on Dataset DPT (see Supplementary Information S2).*

|  | Multiple training | Specific training | Without transfer learning |
| --- | --- | --- | --- |
| Valid loss | 2.406 | 2.416 | 2.56 |
| Number of valid phylogenetic trees | 997 / 1,000 | 998 / 1,000 | 948 / 1,000 |
| Average nRF | 0.087 | 0.099 | 0.513 |

### Supplementary Information S4: Training parameters

Models were trained using mixed-precision arithmetic, in which selected operations are performed in 16-bit rather than 32-bit precision, thereby reducing both memory usage and training time (Micikevicius et al., 2018). We used a cosine learning-rate scheduler, a batch size of 128, and a maximum sequence length of 2,048 tokens. Training included 512 warmup steps and proceeded for up to 20,480 optimization steps. We used 24 layers with a hidden size of 1,024, 16 layers with a hidden size of 1,280, and 12 layers with a hidden size of 1,408, for Configurations 1, 2, and 3, respectively. For each configuration, we tested two learning rates: $1\times{10}^{-4}$ and $1\times{10}^{-5}$.
